# Single-cell characterization of the human C2 dorsal root ganglion recovered from C1-2 arthrodesis surgery: implications for neck pain

**DOI:** 10.1101/2025.03.24.645122

**Authors:** Asta Arendt-Tranholm, Ishwarya Sankaranarayanan, Cathryn Payne, Marisol Mancilla Moreno, Khadijah Mazhar, Natalie Yap, Abby P. Chiu, Allison Barry, Pooja P. Patel, Nikhil N. Inturi, Diana Tavares Ferreira, Anubhav Amin, Mahesh Karandikar, Jeffrey G. Jarvik, Judith A. Turner, Christoph P. Hofstetter, Michele Curatolo, Theodore J. Price

**Affiliations:** Center for Advanced Pain Studies, Department of Neuroscience, University of Texas at Dallas, Richardson, TX, USA; Department of Neurological Surgery, University of Washington, Seattle WA, USA; Department of Anesthesiology and Pain Medicine, University of Washington, Seattle WA, USA; Department of Radiology, University of Washington, Seattle WA, USA; Department of Psychiatry & Behavioral Sciences, University of Washington, Seattle WA, USA; The University of Washington Clinical Learning, Evidence and Research (CLEAR) Center for Musculoskeletal Disorders.

## Abstract

Neurons in the dorsal root ganglion (DRG) receive and transmit sensory information from the tissues they innervate and from the external environment. Upper cervical (C1-C2) DRGs are functionally unique as they receive input from the neck, head, and occipital cranial dura, the latter two of which are also innervated by the trigeminal ganglion (TG). The C2 DRG also plays an important role in neck pain, a common and disabling disorder that is poorly understood. Advanced transcriptomic approaches have significantly improved our ability to characterize RNA expression patterns at single-cell resolution in the DRG and TG, but no previous studies have characterized the C2 DRG. Our aim was to use single-nucleus and spatial transcriptomic approaches to create a molecular map of C2 DRGs from patients undergoing arthrodesis surgery with ganglionectomy. Patients with acute (<3 months) or chronic (≥3 months) neck pain were enrolled and completed patient-reported outcomes and quantitative sensory testing prior to surgery. C2 DRGs were characterized with bulk, single nucleus, and spatial RNA sequencing technologies from 22 patients. Through a comparative analysis to published datasets of the lumbar DRG and TG, neuronal clusters identified in both TG and DRG were identified in the C2 DRG. Therefore, our study definitively characterizes the molecular composition of human C2 neurons and establishes their similarity with unique characteristics of subsets of TG neurons. We identified differentially expressed genes in endothelial, fibroblast and myelinating Schwann cells associated with chronic pain, including *FGFBP2, C8orf34* and *EFNA1* which have been identified in previous genome and transcriptome wide association studies (GWAS/TWAS). Our work establishes an atlas of the human C2 DRG and identifies altered gene expression patterns associated with chronic neck pain. This work establishes a foundation for the exploration of painful disorders in humans affecting the cervical spine.

## Introduction

Since the introduction of single cell and spatial sequencing technologies, a major goal of the somatosensation and pain neuroscience fields has been to characterize dorsal root ganglion (DRG) and trigeminal ganglion (TG) cells with molecular precision (Renthal et al., 2021). Great progress has been made in this area using rodent DRG and TG neurons where we now have comprehensive cell censuses from development to adulthood and including many chronic pain models (Usoskin et al., 2015; Zeisel et al., 2018; Renthal et al., 2020; Sharma et al., 2020). Similar progress has been made in characterizing the human DRG and TG where we also now have spatial and single cell sequencing datasets that demonstrate that there are between 12 and 18 subtypes of sensory neurons in the adult human, lumbar DRG (Nguyen et al., 2021; Tavares-Ferreira et al., 2022; Jung et al., 2023; Bhuiyan et al., 2024; Yu et al., 2024). Subtypes of human TG neurons have also been characterized (Yang et al., 2022), although to a lesser extent than the progress that has been made on lumbar DRGs. These studies have revealed full transcriptomes of neuronal cell types in multiple species and a harmonized atlas across species has recently been built (Bhuiyan et al., 2024). Important species differences have also been found. One major difference between humans and rodents is the more expansive expression of the capsaicin and noxious heat receptor TRPV1 in human DRG neurons (Shiers et al., 2020; Shiers et al., 2021; Tavares-Ferreira et al., 2022). Another is the lack of a clear distinction between peptidergic and non-peptidergic nociceptors in human DRG (Shiers et al., 2020; Shiers et al., 2021) and the identification of unique subsets of human DRG neurons (Tavares-Ferreira et al., 2022; Yu et al., 2024).

A key gap in knowledge is a lack of information about DRGs at the cervical level in humans. These DRGs, at the C1 and C2 level, are functionally and anatomically interesting because they innervate structures like skin, muscle and bone, but also the occipital portion of the cranial meninges (Keller et al., 1985; Noseda et al., 2019). The rest of the cranial meninges is innervated by the TG from the V1 subdivision of that ganglia. Presumably, DRG neurons in the C2 DRG would share characteristics of both DRG and TG neurons, but this hypothesis has never been addressed in humans. Another important gap in knowledge pertains to a common and disabling painful disorder – chronic neck pain. Neck pain is an important cause of disability world-wide; existing treatments are sparse and mostly lack an evidence base for their clinical use (Safiri et al., 2020; Wu et al., 2024). The C2 DRG innervates the atlantoaxial joint, a known originator of neck pain in osteoarthritis, and structures of the cervical spine that can be acutely injured after a fall. In addition, changes to the C2 DRG may be a source of pain. During C1-2 fusion surgeries, this DRG can be resected as part of clinical care.

The work described here had two primary goals. The first was to characterize the transcriptome of cells in the human C2 DRG using single cell and spatial transcriptomic techniques with the purpose of addressing how neurons in these ganglia relate to previous findings in lumbar DRG and TG of humans. The second goal was to ascertain whether the transcriptomes of these cells were altered in samples from people who suffer from chronic neck pain, compared to those with acute pain. This would address the lack of understanding of the transcriptional mechanisms underlying chronic neck pain, potentially identifying novel molecular targets to treat this burdensome condition.

Here we describe the first analysis of the human C2 DRG at single cell resolution. Our work demonstrates that there are similarities to both the lumbar DRG and TG in the human C2 DRG, consistent with its unique anatomical innervation pattern. The removal of these DRGs during C1-2 fusion surgeries, performed to treat either an acute cervical spine injury or chronic neck pain, provided an opportunity to examine changes in gene expression that might be linked to chronic neck pain. We found several endothelial, fibroblast and Schwann cell-related genes that were associated with chronic neck pain, and relatively few changes in neuronal transcriptomes. Altogether, our findings enhance our knowledge of the molecular makeup of human DRGs, expanding our understanding to the cervical level, and give insight into potential mechanisms of chronic neck pain.

## Results

### Characterization of the human C2 DRG with single nucleus sequencing

We used single-nucleus RNA sequencing (snRNA-seq) to characterize the molecular composition of the C2 DRG. DRGs from 12 patients were included in this analysis, 7 DRGs from patients with acute injury and 5 DRGs from patients with chronic pain. Nuclei were isolated from hDRGs and fixed and barcoded using the 10X flex kit to facilitate single-nucleus resolution RNA expression data. We imposed cutoffs of 500 genes and 7.5% mitochondrial-RNA (mtRNA) and obtained a minimum of 129 nuclei per sample. Two rounds of sequencing were performed with more nuclei obtained per sample in the initial run, therefore the RunHarmony tool from the Seurat package was used to merge and integrate the data. Using the FindClusters tool of the Seurat package, we identified 18 unique cell clusters (Fig 1A-B). Expression of neuronal (*SNAP25/PRPH*) and non-neuronal (*MPZ/COL1A1/FABP7*) marker genes correlated with distinct groupings in the UMAP (Fig 1C-D). Representation from all samples was confirmed in all neuronal clusters, however, the non-neuronal cells were largely collected in one run as observed in Figure 1B. We suggest the variability in cell subtype recovery was due to technical variation in the dissociation process. The data was subset into neuronal and neuronal groups, to characterize subclusters in greater detail.

**Figure 1:**
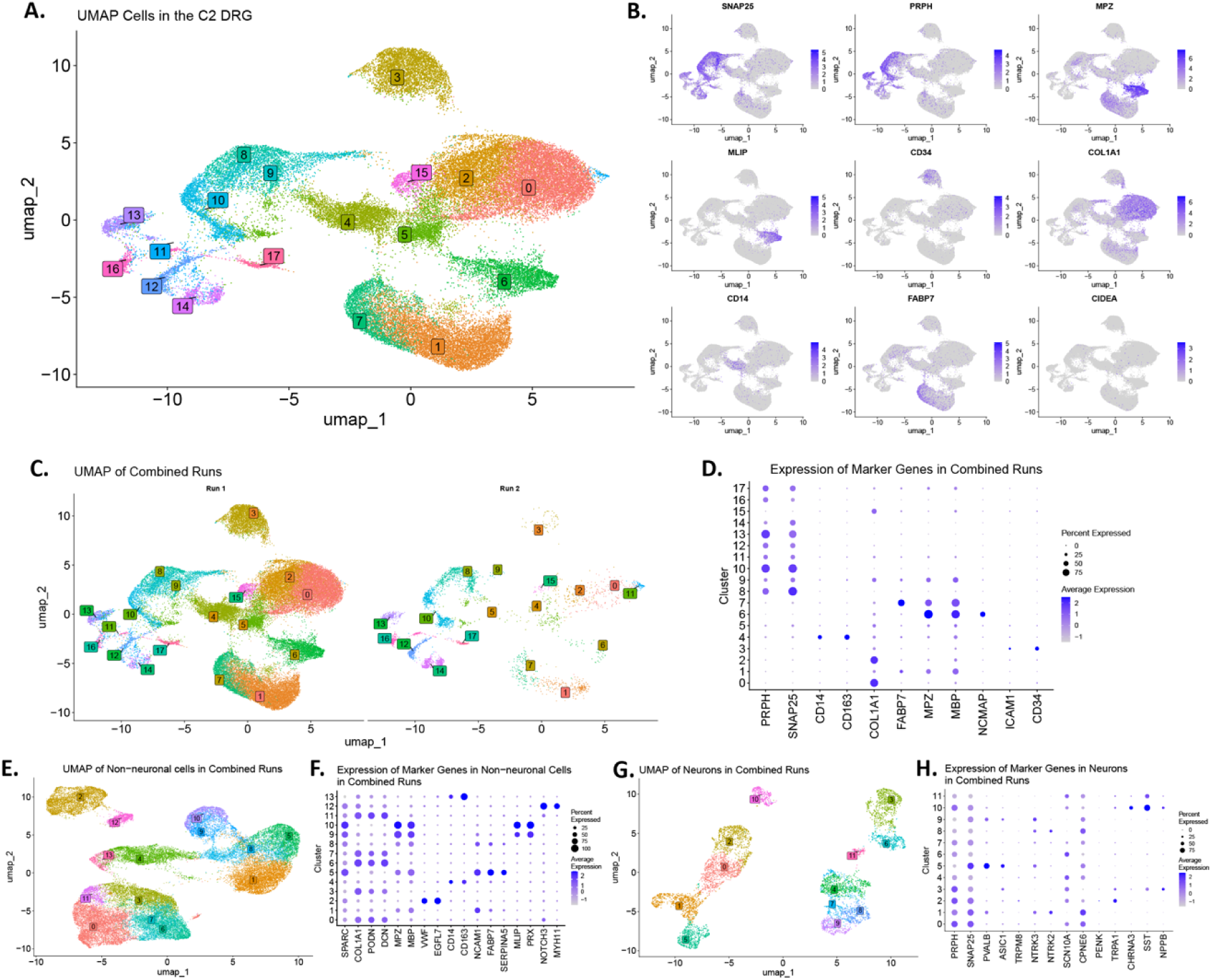
Cell types identified in C2 DRGs using single-nuc RNAseq. **A.** UMAP plot with 18 clusters identified in 12 DRGs from 12 patients. **B.** UMAP plots showing expression of marker genes used to identify neuronal (*SNAP25, PRPH*) and non-neuronal (*MPZ, MLIP, CD34, COL1A1, CD14, FABP7, CIDEA*) cell populations. **C.** Split UMAP showing cell populations identified in 2 runs of single-nuc RNAseq. **D.** Dot plot showing expression of marker genes across the 18 clusters of cells. **E.** UMAP of 14 non-neuronal cell clusters identified by expression of non-neuronal marker genes. **F.** Dot plot of expression of non-neuronal marker genes across 14 non-neuronal clusters. **G.** UMAP of 12 clusters of neuronal cells identified by enrichment of neuronal marker gene expression. **H.** Dot plot of expression of neuronal marker genes across 12 neuronal clusters.

Fourteen non-neuronal cell clusters were identified as seen in Figure 1E. Marker gene expression suggested several of these clusters were of the same non-neuronal cell type (Fig 1F). Expression of *COL1A1, PODN* and *DCN* was distinct in clusters 0, 3, 6, 7, and 11, indicating fibroblast populations. Clusters 4 and 13 exhibited expression of *CD14* and *CD163* indicating immune cell subtypes. Clusters 9 and 10 were identified as Schwann cells based on *MLIP* and *PRX* expression. Clusters 1, 5 and 8 showed variable but distinct expression of *NCAM1* and *FABP7*. Cluster 5 was unique in expressing *SERPINA5* and could be distinguished as a satellite glial cell (SGC) population, while the lack of *SERPINA5* allowed the labeling of clusters 1 and 8 as non-myelinating Schwann cells. Cluster 2 and cluster 12 both showed unique distinct expression of *VWF/EGFL7* and *NOTCH3/MYH11* indicating endothelial and mural cell populations, respectively.

Twelve neuronal clusters were characterized as shown in the UMAP in Figure 1G. Based on marker gene expression clusters 1, 5, 8 and 9 were identified as non-nociceptive (*NTRK2*/*NTRK3*) and clusters 0, 2, 3, 4, 6, 7, 10 and 11 as likely nociceptors (*SCN10A/TRPA1/SST)*), based on marker gene expression (Fig 1H). Additionally, cluster 2 could be identified as TRPA1+ nociceptors, and cluster 10 and 11 expressed *CHRNA3, SST, NNPB* and *OSMR* indicating putative silent nociceptor populations. Other marker genes, including *PENK* and *TRPM8* expressions, did not appear highly expressed. We, consequently, sought to use additional approaches to characterize the neuronal subtypes in greater detail.

### Comparison of C2 DRG neurons to thoracic and lumbar DRG and TG populations

To characterize the neuronal clusters of the C2 DRG, we utilized previously published labeling of human DRG, as well as trigeminal ganglia (TG) (Tavares-Ferreira et al., 2022; Bhuiyan et al., 2024). Using CONOS (Barkas et al., 2019), we transferred labels from neurons characterized in lumbar DRGs onto the C2 DRG data (Fig 2A). The UMAP, which shows a spatial representation of transcriptional similarity between nuclei, showed the groupings identified in C2 DRGs correlating with labels from lumbar DRGs. In Figure 1B, violin plots showed the proportions of clusters from lumbar DRGs identified in each of the C2 clusters.

**Figure 2:**
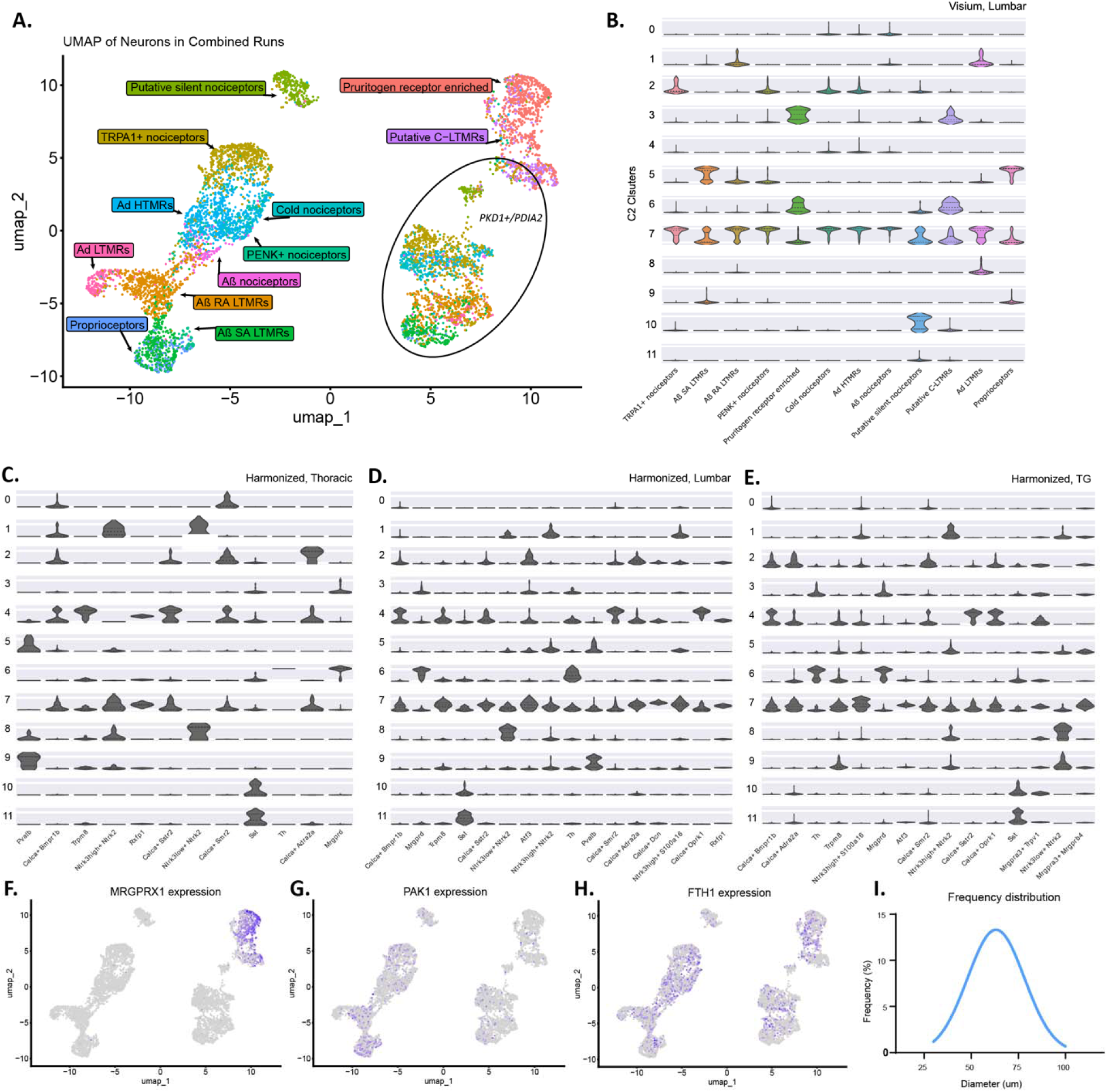
Characterization of neuronal cell subtypes identified in C2 DRG compared to lumbar, thoracic and trigeminal ganglia. **A.** UMAP of neuronal clusters with labels from lumbar DRGs identified with spatial transcriptomics. Cells with enrichment of *PKD1* and *PDIA2* are circled. **B.** Violin plot showing the proportional expression of lumbar DRG subtypes identified with spatial transcriptomics in Tavares-Ferreira & Shiers’ atlas, in each neuronal cluster in the C2 DRG. **C-E.** Violin plot showing the proportional expression of thoracic DRG (**C.**), lumbar DRG (**D.**), or trigeminal ganglia (**E.**) subtypes labeled with single-nuc RNAseq in Bhuiyan et al.’s atlas, in each neuronal cluster in the C2 DRG. **F-H.** UMAP plots showing expression of genes *MRGPRX1* (**F.**), *PAK1* (**G.**) and *FTH1* (**H.**). **I.** Diameters of neurons was measured from H&E-stained images of C2 DRGs and the frequency of diameters was shown (in micrometers). Gaussian curve fits are shown for visualization purposes.

Cluster 1 contained A-beta LTMR and A-delta LTMR neurons. Cluster 2 contained primarily TRPA1+ nociceptors, as well as minor proportions of PENK+ and cold nociceptors. Cluster 10 was unique in distinctly correlating with the labeling of putative silent nociceptors. Interestingly, cluster 7 was found to contain transcriptional similarity to all neuronal subtypes identified in lumbar DRG, while exhibiting a very low number of enriched genes, indicating high transcriptional heterogeneity in the cluster. Enriched expression of *PKD1* and *PDIA2*, two genes that have previously been associated with injured neurons in the study of Parkinson’s disease, was observed in cluster 4, 6, 8, 9 and 11 which surrounded cluster 7 in Figure 1H (Conn et al., 2004; Amadesi et al., 2009; Asaithambi et al., 2011). The probability of transcriptional similarity between clusters 4, 7, 8 and 9 is high, based on the proximal location of nuclei in the UMAP. In fact, the UMAP in Figure 2A with label transfer revealed the clusters in the lower right quadrant appeared to include smaller subsets of all neuronal subtypes, and these clusters all, apart from 7, showed *PKD1* and *PDIA2* enrichment (Supplementary table 1A). Importantly, the source material for this study was patient surgeries, specifically patients receiving surgery for either acute traumatic injury or chronic ongoing degeneration of the joint. Consequently, injured neurons may have originated from the ongoing disease state whether it was chronic neck pain with arthritis or acute neck injury requiring C1-2 fusion. Interestingly, these genes are detected in previous studies of hDRG but are lowly expressed in mouse (Jung et al., 2023; Bhuiyan et al., 2024), and are not known to be changes by injury in the mouse DRG (Renthal et al., 2020), suggesting that these genes could be markers of neuronal injury in C2 hDRG.

We additionally used DRG and TG neuron labels characterized using single-nuc RNAseq in the harmonized atlas by Bhuiyan et al. to assess the similarity to C2 DRG neurons to lumbar and thoracic DRG neurons and TG neurons (Fig 2C-E). *PVALB+* and *DCN+* neuronal populations labeled in lumbar and thoracic DRGs were observed in clusters 5, 7 and 9 in C2 DRGs (Fig 2C-D). *ASIC1* was enriched in clusters 5 and 9 indicating non-nociceptive neurons. Neuronal populations identified in TG defined by *Mrgpra3+* (orthologous to human *MRGPRX* genes) were observed in C2 DRGs, in clusters 2, 3, 4, 5, 7, 8 and 10 (Fig 2E). Additionally, *MRGPRX1* was found to be enriched in clusters 3 and 8 (Fig 2F). *MRGPRX1+* neurons were exclusively identified in TG in human samples in the harmonized atlas (Bhuiyan et al., 2024), however, it was hypothesized that they were a rare cell type in the DRG (Yu et al., 2024). The enrichment of *MRGPRX1*+ neurons may indicate a higher proportion of this cell-type in cervical DRGs, similar to TG (Tavares-Ferreira et al., 2022; Bhuiyan et al., 2024). *TRPM8*, which has previously been shown to be enriched in TG compared to DRG in rodents (Kobayashi et al., 2005), was lowly expressed in C2 DRG, and not enriched in any cluster. *CALCA*, while being expressed in both DRG and TG, has been described as highly expressed in TG and characterized as playing a role in migraine (Goadsby et al., 1990; Ohlsson et al., 2018). *CALCA* was identified as enriched in 4 neuronal clusters (0, 2, 4, 10), correlating with nociceptive clusters.

In mice, characterization of TG in comparison to DRG has revealed distinct transcriptomic patterns. In a study by Megat et al., 381 DEGs were identified as highly enriched in TG. When comparing the genes identified as enriched in TG and DRG in mice and overlapping with enriched genes in each of the C2 clusters, clusters 0, 1, 5, 8 and 9 all had a higher number of enriched genes in the TG (Supplementary table 1B). *Fth1* and *Pak1* were identified as highly enriched in TG. Both of these genes have human orthologues, *FTH1* and *PAK1* (Megat et al., 2019a). *PAK1* was observed to be enriched in clusters 1 and 5 (A-fiber LTMRs and proprioceptors when labeled according to DRG nomenclature) (Fig 2G). *FTH1* was expressed in multiple clusters and not enriched in one specific cluster (Fig 2H). Interestingly, in human lumbar DRGs characterized with spatial sequencing, *FTH1* was observed as broadly expressed in all neuron populations, while *PAK1* was very lowly expressed (Tavares-Ferreira et al., 2022), indicating both species and DRG-level differences in the C2 DRG.

Hematoxylin and eosin (H&E) stained sections of 4 C2 DRGs were used to measure the average neuronal cell size (Fig 2I). The range of diameter of 407 neurons was 30 to 100 µm. This range falls in between the size-ranges reported for neurons in the lumbar DRGs in humans (30-125 µm) (Tavares-Ferreira et al., 2022) and trigeminal neurons (30-80 µm) (Shiers et al., 2023). The mean neuron diameter of C2 DRG neurons was 64 µm, which aligned closely to the lumbar DRG mean of 67 µm, while TG neurons were 41 µm.

In summary, we observed that the neuronal cells in C2 DRGs exhibit similarities to both lumbar DRGs and TG. All neuronal subclusters characterized in lumbar DRGs were identified in C2 DRG, however, physical characterization of C2 DRGs reflected a reduction in size range compared to lumbar DRGs. This finding correlated with the clusters shown to exhibit transcriptional similarities to TGs: Clusters 0, 1 and 5, labeled as A-fiber LTMRs with label transfer and consequently most closely aligned transcriptionally to larger neurons. These A-fiber LTMR clusters also exhibited enrichment of genes characterizing the TG, including *PAK1.* Additionally, clusters 3 and 8, identified as likely pruritogen receptor enriched neurons, are enriched for *MRGPRX1*, a gene showing robust expression in TG and minimal expression in lumbar DRGs.

### Transcriptomic differences between C2 DRGs recovered for acute injury or chronic pain reflect results from transcriptomic/genomic wide association studies

The cervical DRGs were recovered during surgery from patients suffering either acute trauma or long-term osteoarthritis of the joint associated with chronic pain. Based on quantitative sensory testing (QST) and analysis of self-reported pain scores from patients prior to surgery, it is apparent that the chronic pain experience in most of these patients does not match characteristics that would classify the pain of these individuals as having neuropathic pain (Table 1 and 2). Most individuals did not demonstrate signs of mechanical allodynia, and only a small number of patients in the chronic pain group showed signs of cold hypersensitivity. The major difference in terms of QST and pain report measures was the duration of pain, which was much greater in the chronic pain group. This provided a unique opportunity to assess the molecular differences driving chronic neck pain, in a well-characterized subgroup of patients. We analyzed neuronal clusters and non-neuronal clusters, the latter of which were amalgamated into 7 distinct clusters based on marker gene enrichment as described above (Supplementary table 1C), to identify transcriptomic differences in these cell types associated with chronic pain (Fig 3A-B). Differential gene expression was measured with DESeq2 and defined as genes with adj. p-value < 0.05 and |Log_2_FC| > 0.5.

**Figure 3:**
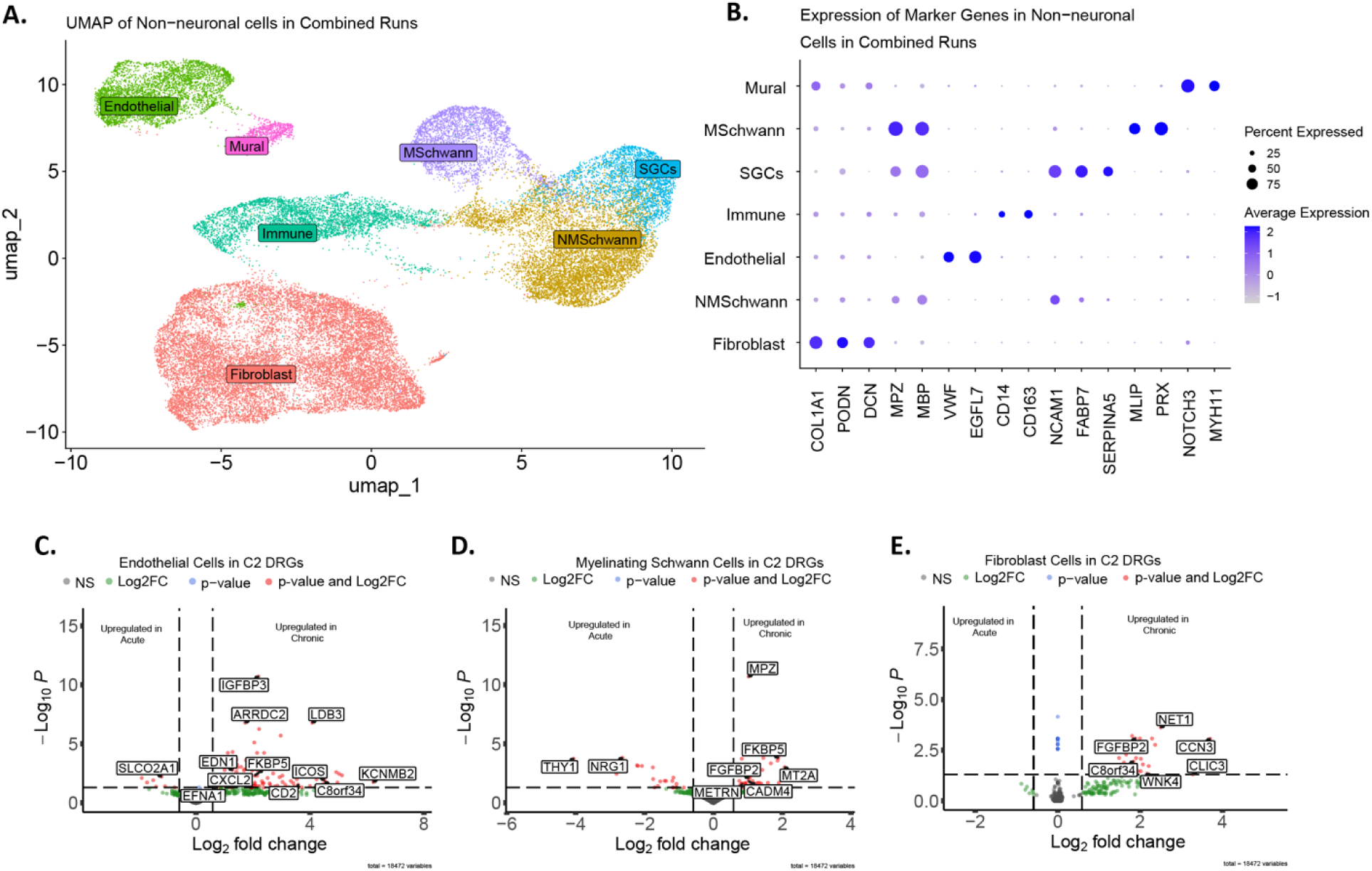
Characterization of non-neuronal cell subtypes identified in C2 DRGs and molecular differences attributed to pain phenotype. **A.** UMAP of non-neuronal clusters labeled through enriched expression of marker genes. **B.** Dot plot showing expression of markers defining non-neuronal cell clusters. **C-E.** Volcano plot showing differential gene expression patterns when comparing samples from acute and chronic patients in endothelial cells (**C.**), myelinating Schwann cells (**D.**) and fibroblast cells (**E.**). Differentially expressed genes (DEGs) were defined as genes with adj. p-value<0.05 and |Log_2_FC|>0.5 and illustrated by red dots.

**Table 1:** Demographic and clinical characteristics.

|  | **All (N= 22)** | **Acute pain (N= 11)** | **Chronic pain (N=11)** |
| --- | --- | --- | --- |
| **Age** |  |  |  |
| Mean (SD) | 71.73 (12.44) | 77.73 (10.74) | 65.73 (11.41) |
| Median (IQR) | 70.50 (63.00-84.00) | 83.00 (64.00-88.00) | 63.00 (57.00-79.00) |
| **Gender, n (%)** |  |  |  |
| Female | 9 (40.91) | 5 (45.45) | 4 (36.36) |
| Male | 13 (59.09) | 6 (54.55) | 7 (63.64) |
| **Body-mass Index (BMI)** |  |  |  |
| Mean (SD) | 28.76 (10.00) | 26.27 (3.57) | 31.25 (13.55) |
| Median (IQR) | 26.54 (23.44-30.53) | 24.41 (23.20-30.22) | 27.18 (23.94-33.73) |
| **Race, n (%)** |  |  |  |
| White | 18 (81.82) | 10 (90.91) | 8 (72.73) |
| Native American/Alaska Native | 1 (4.55) | 0 (0.00) | 1 (9.09) |
| Asian/ Pacific Islander | 2 (9.09) | 1 (9.09) | 1 (9.09) |
| Unknown | 1 (4.55) | 0 (0.00) | 1 (9.09) |
| **Ethnicity, n (%)** |  |  |  |
| Hispanic or Latino | 0 (0.0) | 0 (0.0) | 0 (0.0) |
| Non-Hispanic or Latino | 21 (95.45) | 11 (100.00) | 10 (90.91) |
| Unknown | 1 (4.55) | 0 (0.0) | 1 (9.09) |
| **Use of opioids in past 6 months, n (%)** |  |  |  |
| No | 15 (68.18) | 8 (72.73) | 7 (63.64) |
| Yes | 4 (18.18) | 1 (9.09) | 3 (27.27) |
| Missing | 3 (13.64) | 2 (18.18) | 1 (9.09) |
| **Location of Pain, n (%)** |  |  |  |
| Bilateral pain | 15 (68.18) | 8 (72.73) | 7 (63.64) |
| Unilateral pain | 4 (18.18) | 2 (18.18) | 2 (18.18) |
| Predominantly unilateral pain* | 1 (4.55) | 0 (0.0) | 1 (9.09) |
| Missing | 2 (9.09) | 1 (9.09) | 1 (9.09) |
| **Duration of neck pain, days** |  |  |  |
| Mean (SD) | 193.25 (300.67) | 5.82 (13.37) | 380.68 (335.22) |
| Median (IQR) | 1.00 (1.00-3.00) | 1.00 (1.00-3.00) | 261.00 (169.00-547.50) |
| **Pain in last 24 hours on most painful side\*\*** |  |  |  |
| Mean (SD) | 5.38 (3.27) | 5.75 (3.68) | 5.05 (3.00) |
| Median (IQR) | 6.00 (3.50-8.00) | 6.50 (2.00-9.00) | 4.50 (3.50-8.00) |
| Missing, n (%) | 1 (4.55) | 1 (9.09) | 0 (0.0) |
| **Neck Disability Index (short form)** |  |  |  |
| Mean (SD) | 13.95 (6.36) | 16.78 (6.36) | 11.40 (5.46) |
| Median (IQR) | 13.00 (9.00-21.00) | 20.00 (13.00-21.00) | 12.00 (8.00-14.00) |
| Missing, n (%) | 3 (13.64) | 2 (18.18) | 1 (9.09) |

No differential genes were identified in any neuronal cluster; however, differential expression patterns were discovered in endothelial, fibroblast and myelinating Schwann cells (Fig 3C-E). In the endothelial cluster, 121 genes were identified, 114 of which were upregulated in samples with chronic pain (Fig 3C, Supplementary table 2A). Upregulated genes included *C8orf34*, *ICOS*, *CD2, FKBP5,* and *CXCL2*. *C8orf34* has previously been associated with chronic back pain in multiple GWAS (Freidin et al., 2019; Li et al., 2023; Belonogova et al., 2025). Upregulation of *CXCL2,* a pro-inflammatory chemokine ligand, has been associated with neuropathic pain and orofacial trigeminal pain (Iwasa et al., 2019; Solis-Castro et al., 2021; Hummig et al., 2023; Zeng et al., 2023). Blockage of *FKBP5*, FK506-binding protein 5, has previously been shown to relieve inflammatory pain states in mice (Maiarù et al., 2018). *CD2* is an adhesion molecule usually observed on the surface of T-cells, and associated with an inflammatory state driven by active T-cells. In contrast, *ICOS* has been suggested to alleviate mechanical hypersensitivity induced by paclitaxel through T-cell activation (Sankaranarayanan et al., 2023a). T-cell activity has previously been attributed to both pro- and anti-nociceptive roles depending on the pathway of activation (Ji et al., 2016). Pathway analysis of the DEGs showed enriched Th1 and Th2 cell differentiation as well as IL-17 signaling, implicating endothelial cells in inflammation in chronic pain through interactions with immune cells.

Twenty-five genes were found to be upregulated in fibroblast cells (Fig 3D, Supplementary table 2B). *C8orf34*, *FGFBP2* and *CCN3* were among upregulated genes. *C8orf34* does not have a known function, but was also upregulated in endothelial cells suggesting that it may play a generalized role in non-neuronal cells in arthritic neck pain. *FGFBP2*, which encodes fibroblast growth factor binding protein 2, has been associated with Modic changes in intervertebral disc degeneration and has been implicated as a possible blood biomarker of chronic neuropathic pain (Cherif et al., 2022; Islam et al., 2022; Rajasekaran et al., 2022). *CCN3* was identified as the most highly enriched gene in fibroblasts, and this gene has been identified as playing a key role in the pathology of rheumatoid arthritis through inflammatory fibroblasts (Tokuhiro et al., 2024).

In myelinating Schwann cells, 55 genes were found to be differentially expressed, 33 of which were upregulated in chronic pain (Fig 3E, Supplementary table 2C). *FGFBP2* was upregulated in both fibroblasts and myelinating Schwann cells, again suggesting a generalized role for this gene in arthritic neck pain. *FKBP1*, involved in the stress response and inflammatory pain was upregulated in both Schwann cells and endothelial cells (Maiarù et al., 2018). Furthermore, *MKNK2*, which encodes the mitogen activated protein kinase interacting kinase (MNK) 2, was upregulated. MNK2, is a key mediator of inflammation through translation mediation, and has previously been associated with inflammatory dysregulation driving pain (Moy et al., 2017; Moy et al., 2018). *METRN* and *NRG1* were also identified as differentially expressed. *METRN,* which encodes a secreted protein without a known receptor, was upregulated in chronic pain. Interestingly, previous research suggests a role for *METRN* in resolving neuropathic pain (Jørgensen et al., 2012; Sankaranarayanan et al., 2025). *NRG1*, which encodes neuregulin-1, was upregulated in acute pain. Neuregulin-1 is ascribed a neuroprotective role through immunoregulation driven by the NF-kB pathway (Simmons et al., 2016), however, it has also been indicated to drive neuropathic pain states through phosphorylation of ERK1/2 (Calvo et al., 2011; Dai et al., 2014).

In summary, snRNAseq of C2 DRGs showed the primary transcriptional difference was manifesting in non-neuronal cells. The differentially expressed genes in the endothelial cells, fibroblasts and Schwann cells reflect abnormal transcriptional patterns indicating activation of inflammatory pathways. Simultaneously, multiple transcripts, including *METRN* in Schwann cells and *ICOS* in the endothelial cluster, indicate activation of neuroprotective mechanisms potentially preventing the development of a neuropathic pain state. To assess whether the transcriptional patterns reflected changes to the physical architecture of the DRG we used additional RNA sequencing approaches.

### Inflammatory transcriptomic patterns observed in chronic pain C2 tissues with multiple sequencing approaches

Bulk RNAseq was carried out on 17 DRGs from 13 patients to assess transcriptomic changes in the whole C2 DRG when comparing acute and chronic pain. Fourty DEGs were identified, 17 of which were upregulated and 23 downregulated in chronic pain (Fig 4A, Supplementary table 3A). One gene correlated with findings from snRNAseq, specifically decreased expression of *SLCO2A1*, a transporter of prostaglandin (Fig 4B). The transporter plays a role in clearing prostaglandin E2, PGE_2_, a key mediator of inflammatory pain (Davies et al., 1984; Nakanishi et al., 2021). Additionally, *SLCO2A1* mutations are causative of hypertrophic osteoarthropathy, a disease characterized by inflammation in bones and joints (Lu et al., 2023). Decreased expression of the PGE_2_ transporter suggests a disruption in PGE_2_ metabolism, with the upregulation of PGE_2_ in the C2 DRG contributing to inflammatory dysregulation. Consequently, multiple transcriptional changes are observed in the C2 DRG with bulk RNAseq and snRNAseq which reflect inflammatory pathways that may contribute to abnormal pain signaling. Several genes are observed in more than one cell-type with snRNAseq or in multiple sequencing approaches (Fig 4B-C). To assess the spatial organization of these changes in gene expression, we employed spatial RNAseq.

**Figure 4:**
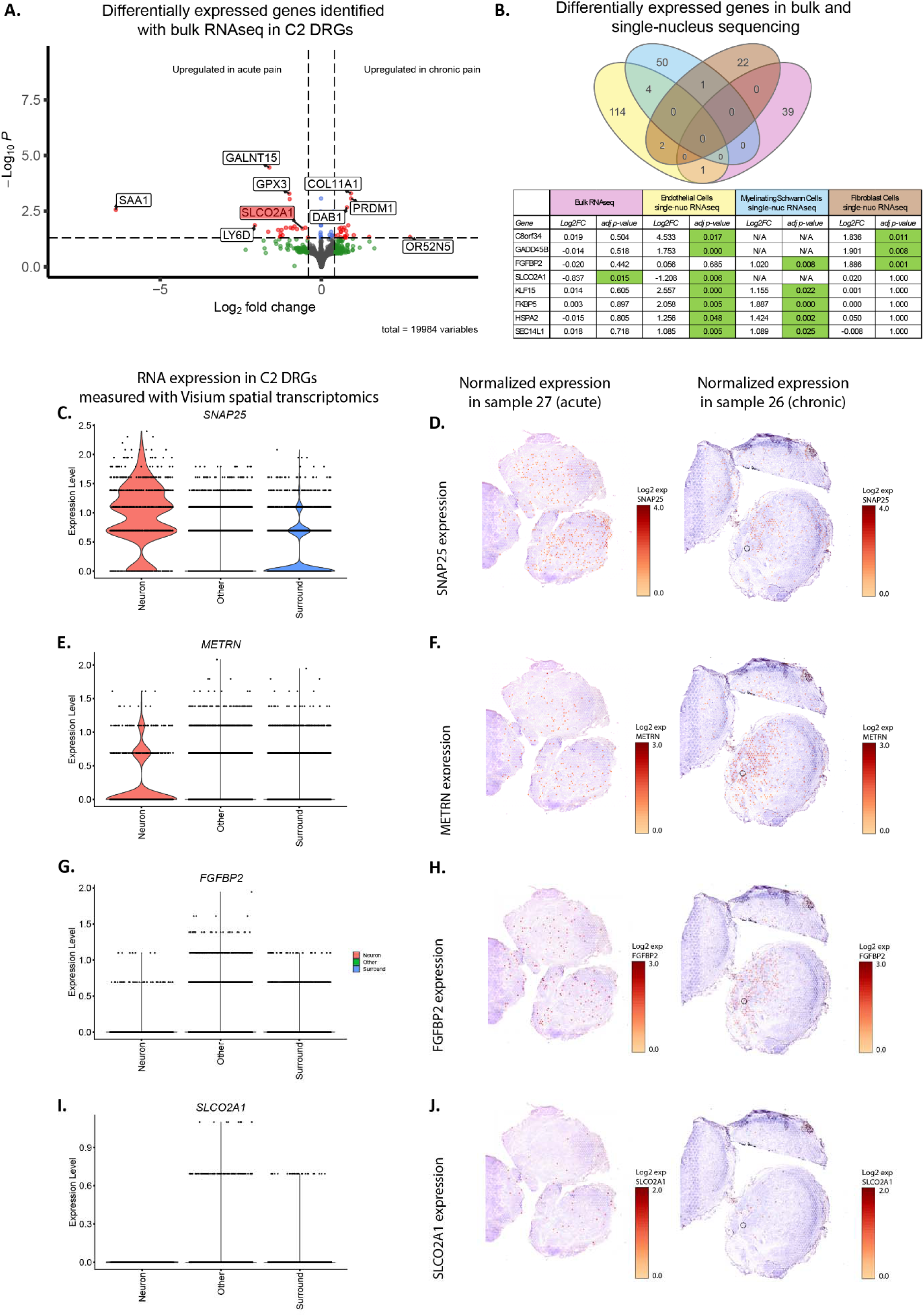
Identifying transcriptome differences in acute and chronic pain samples with bulk and spatial RNAsequencing. **A.** Volcano plot showing differential gene expression patterns identified with bulk RNAsequencing (RNAseq) of 17 DRGs from 13 patients. 40 differentially expressed genes (DEGs) were defined as genes with adj. p-value<0.05 and |Log_2_FC|>0.5 and illustrated by red dots. 17 genes were upregulated in chronic pain and 23 in acute pain including *SLCO2A1*. **B.** Venn diagram showing the overlap of DEGs identified with bulk RNAseq and snRNAseq. Eight DEGs were identified with more than 1 sequencing approach. **C.** Violinplot of expression of SNAP25 identified with Visium spatial RNAseq. SNAP25 is most highly expressed in Neuron barcodes with minor expression in Surrounding barcodes. **D.** Expression of *SNAP25* barcodes shown in red in a sample from a patient with acute pain (27) and a patient with chronic pain (26). **E.** Violinplot of expression of *METRN* identified with Visium spatial RNAseq. *METRN* is most highly expressed in Neuron barcodes. **F.** Expression of *METRN* barcodes shown in red in a sample from a patient with acute pain (27) and a patient with chronic pain (26) near *SNAP25+* barcodes. **G.** Violinplot of expression of *FGFBP2* identified with Visium spatial RNAseq. *FGFBP2* is most highly expressed in Other barcodes. **H.** Expression of *FGFBP2* barcodes shown in red in a sample from a patient with acute pain (27) and a patient with chronic pain (26) near *SNAP25+* barcodes. **I.** Violinplot of expression of *SLCO2A1* identified with Visium spatial RNAseq. *SLCO2A1* is most highly expressed in Other barcodes. **J.** Expression of *SLCO2A1* barcodes shown in red in a sample from a patient with acute pain (27) and a patient with chronic pain (26).

We obtained spatial transcriptomic data from 13 C2 DRGs from 8 patients. Sections of DRG were sectioned onto Visium slides with 55 µm barcoded spots and imaged following H&E staining to obtain spatial information. Unique barcodes corresponding to spatial location were bound to mRNA in each section to obtain spatially correlated transcriptomic data. We visually confirmed barcodes that overlapped neurons and defined barcodes as (1) neuronal, (2) immediately surrounding a neuron (surrounding), and (3) others, as defined previously (Tavares-Ferreira et al., 2022). No differentially expressed genes were identified when comparing samples from acute and chronic patients in any group of barcodes. This may be due to insufficient sample size to find differences using this technique. However, the Visium data allowed for a spatial characterization of the DEGs from bulk- and snRNAseq. The expression of *SNAP25* was primarily observed in neuronal barcodes, validating the barcode groups (Fig 4D). *SNAP25* consequently visualizes the neurons in samples 26 (chronic pain) and 27 (acute pain) in Figure 4E-F. Subsequently, we evaluated the spatial expression patterns of *FGFBP2*, *METRN* and *SLCO2A1*. *FGFBP2* was upregulated in chronic pain in myelinating Schwann cells and fibroblast cells, while *METRN* was upregulated in Schwann cells. Using the spatial transcriptomics data, *FGFBP2* and *METRN* expression in sample 26 appears similar, visually localizing near the areas with *SNAP25* expression. The remaining *SLCO2A1+* barcodes were also proximal to *SNAP25*+ areas. However, when considering all samples, *METRN* was primarily measured within neuronal barcodes, while *FGFBP2* was measured in Other barcodes (Fig 4). In contrast, the expression of *SLCO2A1*, which was downregulated in chronic pain with bulk RNAseq and snRNAseq, was absent in neuronal barcodes. The expected consequence of this downregulation would be increased PGE_2_ concentration in the area around C2 DRG neurons.

### Predicted communication between neuronal and non-neuronal cells driving chronic pain states

The transcriptional changes in the chronic pain C2 DRG indicate a reorganization of the transcriptomes of certain non-neuronal cell types with little change in the neuronal populations. In order to assess how the non-neuronal cells may exert an impact on neuronal cells, we used interactome analysis. We assessed ligand genes that were upregulated in non-neuronal cell types in chronic pain C2 DRGs and intersected these genes with receptors for these ligands that are expressed by different populations of C2 DRG neurons. This analysis showed interactions between non-neuronal (endothelial, fibroblast and myelinating Schwann cells) and neuronal cells corresponding to a curated database of ligand-receptor interactions (Fig 5A-C). *EDN1* and *EFNA1* were identified as expressed in endothelial cells with multiple receptors and interactions in neurons: *EFNA1*, which encodes an ephrin protein, interacted with tyrosine kinase receptors encoded by *EPHA* and *EPHB* genes (Fig 5A-B). *EPHA3/5* showed higher expression in nociceptor cells, indicating a more likely interaction with nociceptors, while the remaining *EPHA* genes were more highly expressed in non-nociceptive neuronal clusters (Fig 5C). *EDN1* interacts with *EDNRA*, *KEL* and *ADGRL4*. *EDN1* interaction with the receptor *EDNRA* is likely the most important in this context as a genome association also study found correlation between *EDNRA* and migraine (Lisi et al., 2006; Miao et al., 2012), previous studies have shown that endothelin 1 causes pain like behavior in rodents through the *EDNRA* receptor (Zhou et al., 2001; Zhou et al., 2002), and the *EDN1-EDNRA/EDNRB* axis has been linked to oral cancer pain (Zhou et al., 2002; Dang et al., 2020). *CSF3* and *IL34* were upregulated in chronic pain in endothelial cells, and found to interact with *CSF1R* and *CSF3R*, both of which showed higher expression in nociceptor subtypes (Fig 5B-C). *IL34* encodes a cytokine that interacts with *CSF1R*, which has previously been associated with inflammation, as well as several pain conditions including bone pain, arthritis, and neuropathic pain (Schweizerhof et al., 2009; Bali et al., 2013; Guan et al., 2016; Ushach and Zlotnik, 2016; Lee et al., 2017; Lin et al., 2019) Additionally, *CXCL2*, a chemokine characterized in neuropathic pain, and specifically in trigeminal pain, was found to interact with three chemokine receptors in neurons (Iwasa et al., 2019; Solis-Castro et al., 2021; Hummig et al., 2023; Zeng et al., 2023).

**Figure 5:**
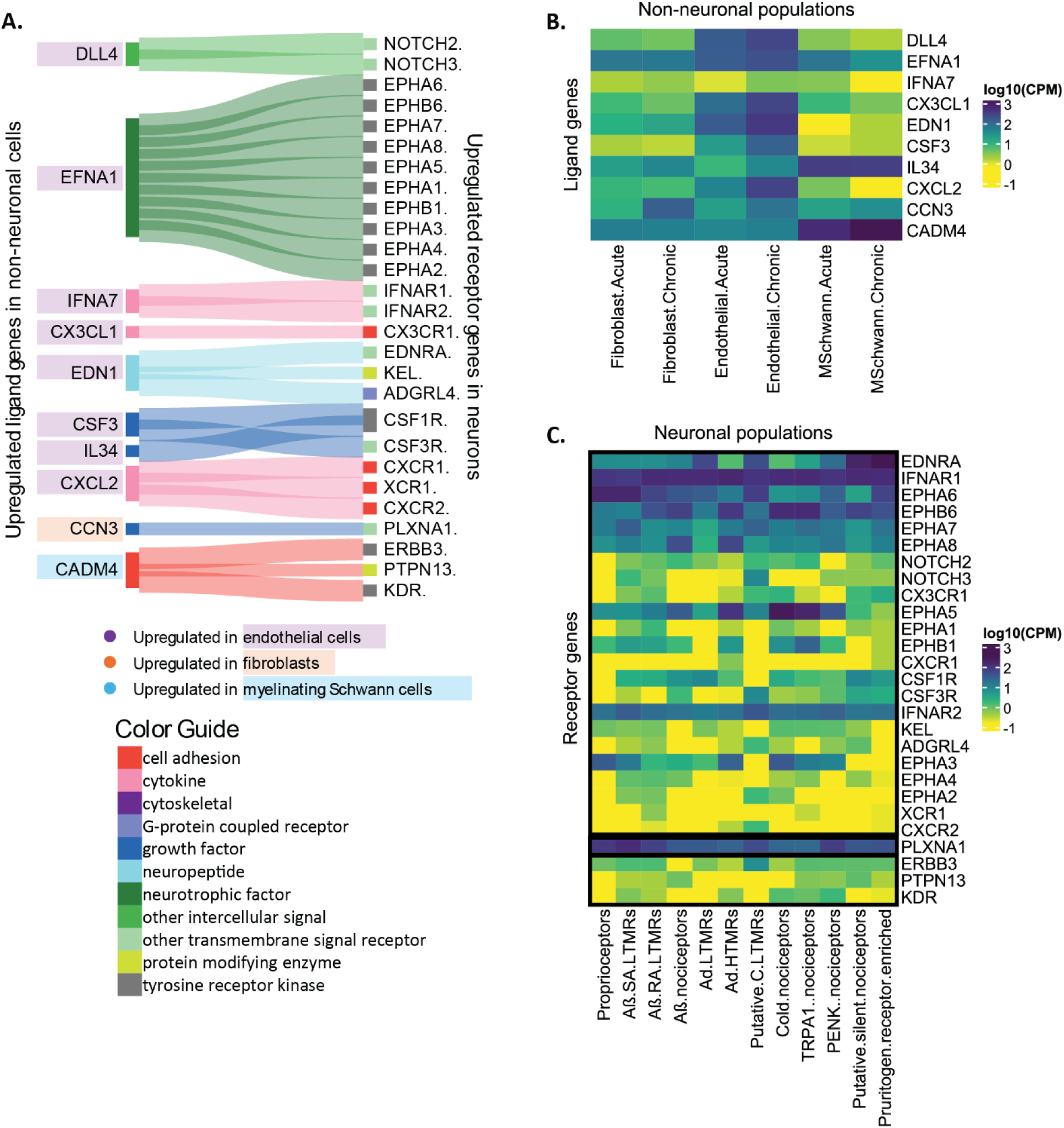
Interactome analysis showing interactions between ligands released from non-neuronal cells in chronic pain states and receptors in neurons. **A.** Sankey diagram showing predicted interactions between ligands released from non-neuronal cells in chronic pain states (defined as genes with adj. p-value<0.1 and |Log_2_FC|>0.5) and receptors expressed in neurons. **B.** Heatmap showing the expression of ligands expressed in non-neuronal cells in acute and chronic pain samples. **C.** Heatmap showing the expression of receptors expressed in neuronal cells.

Fewer differentially expressed genes were identified in myelinating Schwann cells and fibroblasts, consequently the identified interactions with neurons were limited (Fig 5A-B). *CCN3*, which was upregulated in fibroblasts and previously characterized in arthritis (Tokuhiro et al., 2024), was indicated to interact with *PLXNA1*, which was broadly expressed in neurons (Fig 5A-C). *CADM4* is a cell adhesion molecule, the interactions of which exhibited a trend towards increases in nociceptors (Fig 5C). *CADM4* activity with *ERBB3* and *PTPN13* has primarily been characterized in relation to myelin formation, and an increase in CADM4 expression has been associated with abnormal myelination formation (Elazar et al., 2019). *CADM4* additionally interacts with *KDR*, which encodes the receptor VEGFR2. VEGFR2 has been associated with synapse formation and neuronal guidance (De Rossi et al., 2016). The dysregulation of CADM4 could consequently be theorized to impact innervation.

Altogether, a transcriptome pattern reflecting the upregulation of regulators of inflammation, mediated by non-neuronal cells, is observed in C2 DRGs from patients with chronic pain. The transcription patterns are reflected both in the proximal area of neurons as well as in distant areas, suggesting widespread dysregulation across neuronal and non-neuronal networks.

## Discussion

Our work on C2 DRGs recovered from patients undergoing arthrodesis surgery with ganglionectomy led to the following major conclusion. First, consistent with the innervation patterns for C2 DRGs that share properties with lumbar DRGs and TGs, we found that these neurons represent a mixed transcriptomic phenotype between DRG and TG neurons. Second, we found that most transcriptomic changes in chronic pain C2 DRGs were associated with non-neuronal cells including endothelial, myelinating Schwann cells, and fibroblasts. These changes were mostly associated with inflammation and suggest that arthritis of the atlantoaxial joint in humans causes inflammatory-related changes in gene expression in the associated C2 DRG. Finally, integrating our single nucleus sequencing results with spatial transcriptomic data and a ligand-receptor interaction database revealed potential mechanisms that might be pain generators within the C2 DRG in individuals with chronic neck pain. These mechanisms included signaling molecules like endothelin-1 (Zhou et al., 2001; Zhou et al., 2002), FKBP5 (Linnstaedt et al., 2018; Geranton, 2019; Maiaru et al., 2023; Ahlstrom et al., 2024; Lafta et al., 2024), and MNK2 (Moy et al., 2017; Moy et al., 2018; Megat et al., 2019b; Shiers et al., 2023; David et al., 2024; Li et al., 2024; Mitchell et al., 2024) that have previously been associated with pain in animal models and human studies, and mechanisms that might promote neuronal sprouting which could cause pathological changes in innervation in chronic neck pain.

The human C2 DRG appears to share properties of more caudal DRGs and the TG. The C2 DRG is unique because it innervates areas of the upper neck but also contain neurons that innervate the occipital dura, a specialized tissue protecting the brain that is also innervated by the TG and implicated in migraine and other headache disorders (Keller et al., 1985; Noseda et al., 2019). This anatomical connection is one of the explanation why painful lesions of the upper cervical spine can be associated with headache. (Bogduk and Govind, 2009). We found several genes that were highly expressed in the C2 DRG that are also enriched in the TG, such as *FTH1* and *PAK1* which were identified in a study in mice (Megat et al., 2019a), and MRGPRX1 which was identified in humans as enriched in the TG (Bhuiyan et al., 2024). While many different atlases of hDRG neurons have been published to-date (Nguyen et al., 2021; Tavares-Ferreira et al., 2022; Jung et al., 2023; Bhuiyan et al., 2024; Yu et al., 2024), none of them have focused on understanding variation in gene expression based on the level of the DRG. Our work shows that the C2 DRG has some unique transcriptomic features and makes the case that future work should strive to better understand molecular features of human sensory neurons along the entire caudal to rostral extent of the peripheral nervous system from sacral DRGs to the TG.

We did not find evidence of neuropathic pain in our chronic neck pain cohort in somatosensory measures, except cold allodynia in 2 patients (Table 2), or in the transcriptome of the C2 DRGs of these individuals. Neck pain is considered of primary musculoskeletal nature, but neuropathic components are possible. The C2 DRG is located between the arch of C1 and the lamina of C2. In this space, the DRG occupies 76% of the foramen height, rendering the ganglion vulnerable to compression in patients with altered spine due to arthritis or fractures (Lu and Ebraheim, 1998). Recent studies of patients with whiplash injury suggest that there is a prominent neuropathic phenotype in these patients with loss of sensory function, mechanical and cold hypersensitivity, neuroinflammation evident with imaging, and increased measures of neurofilaments in plasma, consistent with neuronal injury (Fundaun et al., 2025; Ridehalgh et al., 2025). Table 2 shows some more neuropathic pain characteristics in the acute pain cohort, which may be closer to the whiplash injury phenotype.

**Table 2:**
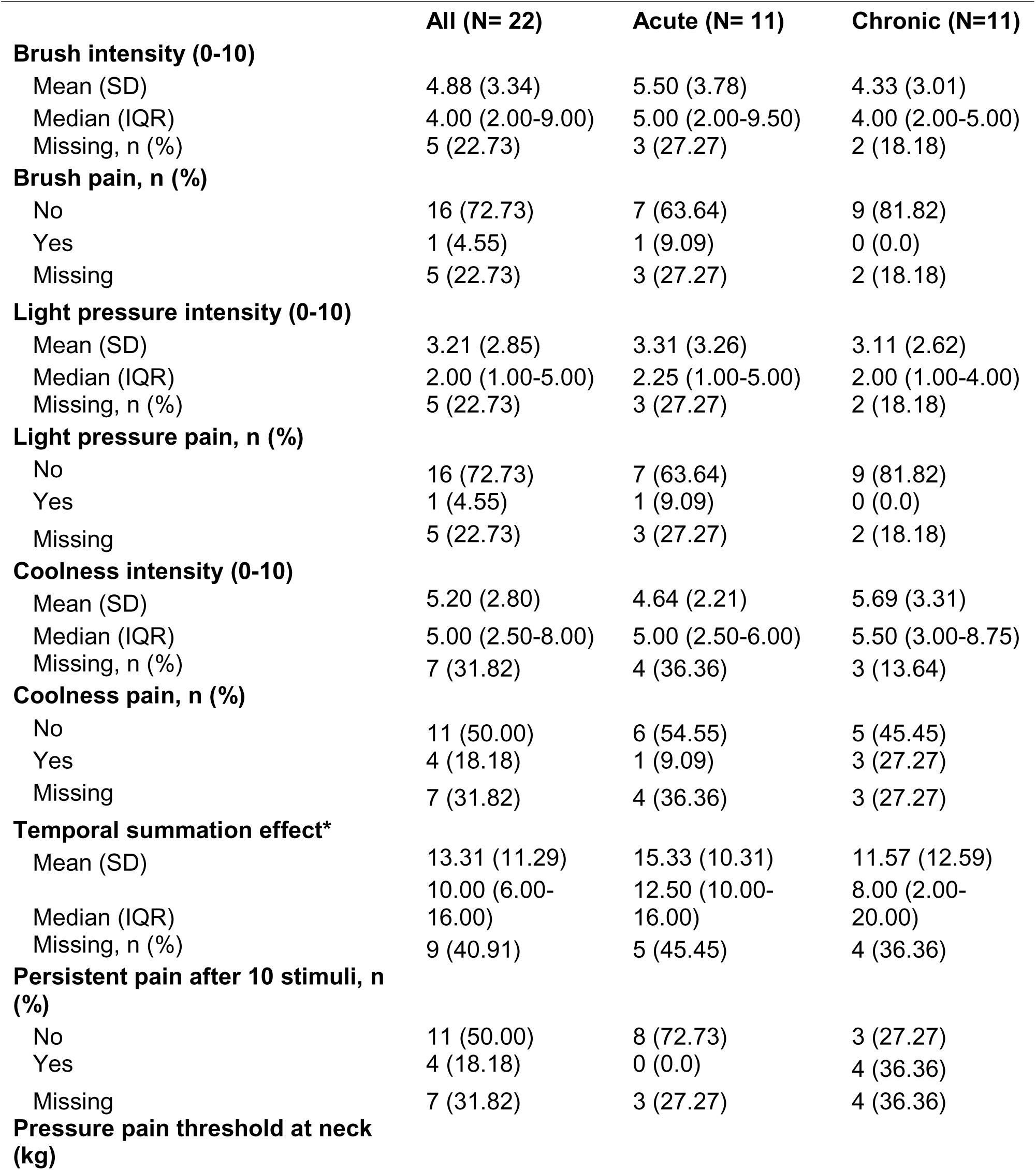
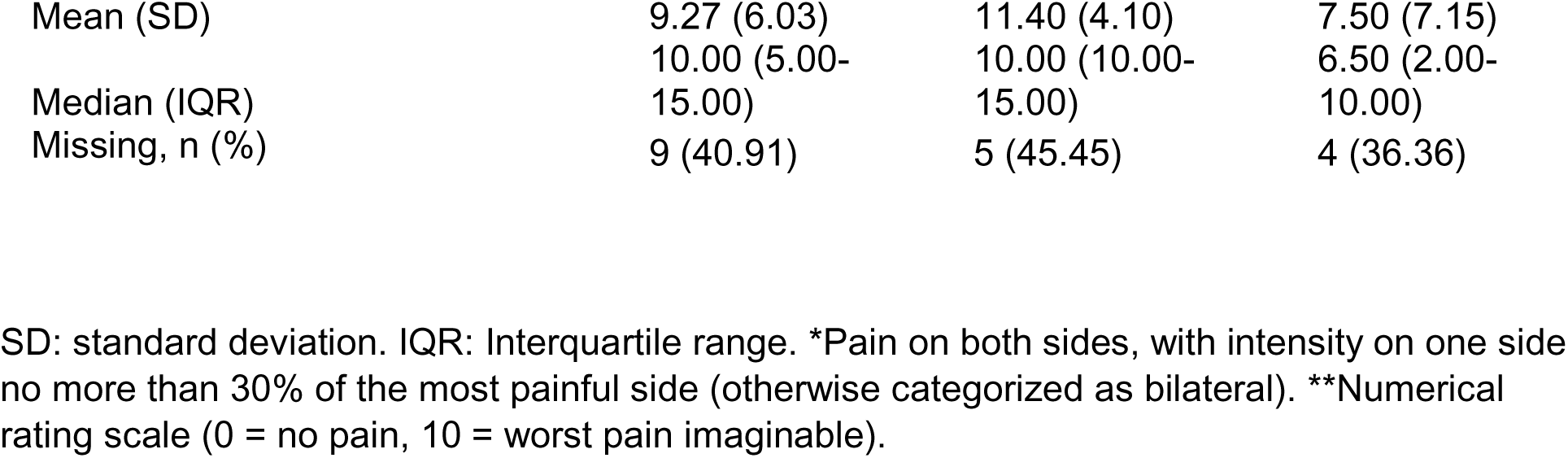
Somatosensory measures.

In DRGs from chronic pain patients in our study, we noted some changes in the neuronal transcriptome that could be associated with neuronal injury (in both the acute and chronic cohort) that were characterized by neurons expressing *PKD1* and *PDIA2*. These neurons mostly had similar transcriptomes to matching populations within the C2 DRG not expressing these genes, and did not undergo wholesale changes in their transcriptomes as has been reported in animal models of nerve injury (Renthal et al., 2020). To our knowledge, *PKD1* and *PDIA2* have not previously been reported as markers of nerve injury in the DRG, but they have been associated with nerve injury in Parkinsons disease (Conn et al., 2004; Amadesi et al., 2009; Asaithambi et al., 2011). A question that emerges from these findings is that if there is evidence of nerve injury, and neuropathic pain is common among people with chronic neck pain, why do individuals in this cohort fail to develop signs of neuropathic pain? One possibility is that mechanisms that suppress neuropathic pain are also induced in the DRG of these individuals.

For instance, we observed increased *ICOS* and *METRN* expression in single nuc RNA sequencing data in patients in the chronic pain cohort. Previous work suggests that *ICOS* stimulates T-cells to produce interleukin-10 to suppress neuropathic pain (Sankaranarayanan et al., 2023a). Several studies have shown that meteorin, the protein encoded by the *METRN* gene, can reverse established neuropathic pain states in both mice and rats (Jorgensen et al., 2012; Xie et al., 2019; Sankaranarayanan et al., 2023b). While this idea clearly requires further exploration, it is interesting that pain promoting and pain-resolution mechanisms seem to be induced at the same time in the DRGs of the individuals with chronic neck pain studied here. Another explanation for the limited evidence for neuropathic components is our limited ability to detect painful neuropathy, which relies mostly on somatosensory measures at the skin that may not capture painful neural lesions of deep tissues.

Our study identifies potential mechanisms of chronic neck pain that can be targeted by therapeutics. These include *FKBP5*, endothelin-1 and the endothelin A receptor, MNK2 signaling, and *IL34* acting via the *CSF1* receptor. All of these signaling factors have previously been implicated in pain, but none of them have been associated with neck pain, to our knowledge. This is not surprising because there are few animal models of neck pain and the disorder, while both prevalent and disabling in the human population, is challenging to study at the molecular level. Therefore, an important strength of this study is the molecular insight our work might give to developing new targets for the treatment of neck pain. *FKBP5* encodes the FKBP51 protein that is involved in glucocorticoid signaling. The protein has been implicated in the important link between stress and pain and could be targeted with inhibitors of the protein (Linnstaedt et al., 2018; Geranton, 2019; Maiaru et al., 2023). We identified increased endothelin-1 gene expression in endothelial cells of the chronic pain cohort. The receptor for this protein, the endothelin A receptor, is highly expressed in rodent and human nociceptors and activating the receptor causes pain in rodents through signaling cascades that involve voltage gated sodium channels (Zhou et al., 2001; Zhou et al., 2002; Bhuiyan et al., 2024). *MNK2* encodes a mitogen activated protein kinase (MKNK2) that has been implicated in pain through mouse genetic studies and studies on hDRG neurons. Inhibiting MNK with the specific inhibitor eFT508 leads to decreased pain in mouse studies and a reduction in excitability in human nociceptors (Moy et al., 2017; Megat et al., 2019b; Li et al., 2024). IL34 was upregulated in endothelial cells in the DRG of chronic pain patients in this cohort. The receptor for this cytokine, CSF1R is strongly expressed across neuronal populations in the human C2 DRG but the consequences of activation of CSF1R in hDRG have not yet been studied. We propose that these targets should be prioritized for future investigation as therapeutic interventions for non-neuropathic chronic neck pain.

### Limitations

This study was designed to identify changes in a unique subpopulation of patients, by utilizing an opportunity to recover tissues otherwise discarded during surgery. However, as the primary source was surgical tissue, we did not have access to control samples from a patient population without pain. The physical location of the cervical DRGs in the upper neck makes it challenging to access them during a standard organ donor tissue recovery without an additional incision that can interfere with subsequent tissue recovery for tissue donation. Since recovery of DRGs following organ donation has become an important source of these tissues for research purposes (Shiers et al., 2024), this issue will need to be overcome to obtain control samples of C2 DRGs.

Another limitation is the small sample size in this study. C1-2 arthrodesis is an uncommon procedures, and resection of the DRG as part of clinical care is not uniform practice among surgeons. Given the difficulty in obtaining these samples, and the lack of information on C2 DRG in humans we think that the insight provided from this study justifies the small sample size. Previous work on thoracic DRGs recovered during thoracic vertebrectomy surgery demonstrated that findings from an initial, smaller cohort, were replicated in a larger cohort collected over a longer period of time (North et al., 2019; Ray et al., 2023). We plan to use a similar approach as this unique cohort of patients continues to grow.

## Materials and methods

### Study design

The goal of this study was to characterize human C2 DRGs with molecular precision. For this purpose, DRGs were collected from patients during C1-C2 arthrodesis surgery with ganglionectomy for acute or chronic pain. All human tissue procurement protocols were reviewed and approved by the University of Texas at Dallas (UTD) and University of Washington (UW) Institutional Review Boards. Patient information, including demographics and characterization of clinical presentation, was collected from medical records and self-reported questionnaires prior to surgery.

### Participants and setting

Patients eighteen years or older undergoing C1-C2 arthrodesis for neck pain at the University of Washington were prospectively recruited. The study was reviewed and approved by the University of Washington Internal Review Board (IRB number 10916). Informed consent was obtained from all patients, who were offered a $125 gift card for participating in the study.

Patients were identified from emergency care settings and clinic visits. We screened for patients who were greater than eighteen years old, undergoing C1-C2 arthrodesis. These patients were identified in a two-pronged manner, whereby the surgeon would notify the research staff of scheduled procedures, and by daily electronic reports accessing the feeds of surgery scheduling data, using the UW Medicine Enterprise Data Warehouse. Exclusion criteria were cognitive deficits or language comprehension that would make the patient unable to consent. Potentially eligible outpatients were contacted via phone call to set up a time to meet in person and inpatients were met in their hospital room if they indicated interest in potentially participating in the study. The research staff then informed the participant of the study and consent the patients using a standardized consenting script.

Participants were categorized as having acute (lasting <3 months) and chronic pain (≥3 months), in keeping with the definitions of the International Association for the Study of Pain (IASP) (Treede et al., 2019).

### Sociodemographic variables and patient-reported outcomes

Age, sex, body-mass index (BMI), opioid medication, race, and ethnicity were recorded within two weeks of the surgery from the electronic health records (EHR) when available, or were otherwise confirmed by patient interview.

Measures of pain included location of pain, duration of pain (days), average pain in the last 24 hours on the most painful side of the neck, and neck-related function. Patients were asked to rate their average neck/occipital pain during the last 24 hours on their most painful side using the numerical rating score (NRS), with 0 = no pain and 10 = worst pain imaginable (Dworkin et al., 2005). Patients were asked to indicate the duration of their pain, which was recorded in days. Neck-related function was assessed using the 5-item version of the Neck Disability Index (NDI-5) (Walton and MacDermid, 2013).

### Somatosensory measures

Patients underwent quantitative sensory tests (QST) according to published guidelines (Cruccu et al., 2010) using a standardized and widely used protocol (Backonja et al., 2013). The QST was performed on the most painful side of the neck (or randomly selected side in case of equal bilateral pain). The instruments involved include a soft brush to simulate light brushing, a monofilament to simulate light pressure, and a tuning fork that has been in ice water to simulate coolness. Patients were asked to rate the sensation of brushing, pressure, and coolness from a 0 = felt nothing to 10 = extremely intense feeling, and secondarily whether any of the three modalities cause pain.

Temporal summation, the increased perception of pain in response to sequential stimuli of equal physical strength, was used as an indicator of facilitated pain modulation processes at the neck (Price, 1972; Arendt-Nielsen et al., 1994; Staud et al., 2007). A pinprick stimulator was used to measure the intensity of a single point of stimulus using a 0-100 NRS (0 = no pain, 100 = worst pain imaginable). Ten additional stimuli were subsequently administered, and patients were asked to rate that sensation using the same scale. The temporal summation effect was calculated by subtracting the pain intensity after 10 stimuli by the pain intensity after one stimulus.

Pressure pain sensitivity was conducted using a pressure simulation device (Wagner FDK 40). Gradually increasing pressure at the rate of (0.5 kg-force/s) was administered, and the participant was instructed to inform the researcher of the moment that the sensation changes from a feeling of pressure to a feeling of pain by voice (e.g., ‘stop’, ‘now,’). The recorded pressure (kg) was defined as pressure pain threshold.

### Tissue collection

DRGs were collected from 11 patients with chronic pain and 11 patients with acute pain according to previously described protocol(Hofstetter, 2025). Briefly, C1 lateral mass screws were placed at the midpoint C1 posterior arch and the lateral mass, and C2 pars screws were placed at a parallel trajectory to the C2 pars to ensure fixation. A bilateral C2 DRG neurectomy was performed and DRGs were immediately placed in powdered dry ice to ensure uniform freezing to preserve RNA. DRGs were shipped to UTD for all sequencing experiments.

### Single-nucleus RNA sequencing

To achieve characterization at a molecular level, we used 10X single-cell RNA-sequencing of 12 DRGs from 12 patients. Briefly, DRGs were thawed in artificial cerebrospinal fluid (aCSF) and perineurium as well as cauterized exterior tissue was removed to expose clean C2 DRG bulb.

The bulb was dissociated and filtered to minimize myelin contamination and obtain individual nuclei according to previously published 10X protocol (Lacagnina et al., 2024). The RNA from each nucleus was barcoded and libraries were generated according to the manufacturer’s protocol for sequencing on a NextSeq2000 at the genomic core at the University of Texas Dallas or Psomogen.

### Spatial RNA sequencing

We used 10X Visium spatial transcriptomics to achieve near-single resolution data, retaining concurrent transcriptional and spatial information, according to previously published protocol (Tavares-Ferreira et al., 2022). Briefly, 13 DRGs from 8 donors were sectioned onto Visium slides, stained with hematoxylin and eosin (H&E) and imaged. Tissue permeabilization and binding of oligo primers were performed according to the manufacturer’s instructions.

Sequencing of mRNA libraries was performed on a NextSeq2000 at the genomic core at the University of Texas Dallas and barcodes overlapping neurons were identified visually and confirmed through expression of neuronal marker *SNAP25*.

### Bulk RNA sequencing

To assess transcriptomic differences to the whole DRG associated with pain phenotype, we used bulk RNA sequencing. Briefly, RNA was extracted from 23 C2 DRGs from 13 donors. Three 100µm sections from the bulb were lyzed in TRIZol and RNA extracted with chloroform. DNA was depleted using Qiagen DNase I, according to manufacturer’s instructions. RNA integrity was assessed with nanodrop and Qubit and samples with RNA integrity number 2.4-8.1 were sequenced using NextSeq2000 for paired-end mRNA sequencing. Five samples were excluded due to low gene expression of neuronal markers according to a previous analysis (Ray et al., 2023), thus 17 DRGs from 13 donors were included in downstream analysis.

### Computational analysis

#### Single-nucleus RNA sequencing Pipeline

Sequencing data were processed and mapped to GRCh38 human reference genome (GENCODE release 38) according to 10X protocol using package cellranger. Nuclei with less than 500 genes or more than 7.5% mtRNA were excluded from downstream analysis. Unbiased clustering was performed using the Seurat package v5.2.1. Neuronal and non-neuronal clusters were identified through marker gene enrichment of neuronal markers (*SNAP25*, *RBFOX3* and *PRPH*) and non-neuronal markers (*SPARC, MPZ* and *DCN*). FindAllMarkers was used to identify gene enrichment (|Log_2_FC>0.5|, adj. p-value<0.05 and min. percent expression in 25% of cluster). The neuronal and non-neuronal clusters were subset into separate objects for downstream analysis. Counts were aggregated for cells of the same cluster for each donor, for differential gene expression analysis with DESeq2 (Love et al., 2014). Differential gene expression analysis was carried out to account for batch effects and sex differences: design = ∼ Sex + Batch + Condition. Differentially expressed genes (DEGs) were defined as |Log_2_FC>0.5|, adj. p-value<0.05.

#### Bulk RNA sequencing Pipeline

Raw sequencing (FASTQ) files were processed with FastQC v0.11.0 to examine Phred scores, per-base sequence, and duplication levels. Reads were soft-clipped (12 bases per read) to discard adapters and low-quality bases during alignment using STAR v2.7.6 (Dobin and Gingeras, 2015) with the GRCh38 human reference genome (GENCODE release 38, primary assembly). Reads were also sorted with STAR and then deduplicated with sambamba v0.8.2 (Tarasov et al., 2015). Genes in the ENCODE blacklist were filtered out prior to quantification using bedtools (v2.30.0), and StringTie v2.2.1 was used to obtain normalized count values as transcripts per million (TPM) for each gene of all samples (Quinlan, 2014; Pertea et al., 2016; Amemiya et al., 2019). The featureCounts function of the R package Rsubread (v2.14.2) was used to calculate raw counts for all genes, which was subsequently used for differential expression analysis with DESeq2 (Love et al., 2014; Liao et al., 2019). Differential gene expression analysis was carried out to account for sex differences: design = ∼ Sex + Condition. Differentially expressed genes (DEGs) were defined as |Log_2_FC>0.5|, adj. p-value<0.05.

#### Spatial RNA sequencing Pipeline

Spatial RNA sequencing data was analyzed according to previously published 10X protocols. Briefly, FASTQ files were obtained from the 10X Space Ranger’s mkfastq pipeline. Space Ranger count pipeline was used align H&E-images and barcode count information and reference transcriptome (GENCODE release 38). Neuron, Surrounding and Other barcodes were identified as previously described (Tavares-Ferreira et al., 2022). Cloupe files were obtained to illustrate gene expression in a spatial context. Differential gene expression analysis was carried out with DESeq2 as described in Single-Nucleus RNA sequencing pipeline. Neuron cell diameter was measured from H&E images in CellSens.

#### CONOS cell subtype predictions

Label propagation from previously published lumbar data to the C2 data was performed with the R package conos v1.5.2 (Barkas et al., 2019). A conos object was built on the corresponding, processed RNA assays with basicSeuratProc, using umap = TRUE. A graph was then built in CCA space with the following parameters: con$buildGraph(k=30, k.self=5, space=’CCA’, ncomps=40, n.odgenes=2000, matching.method=’mNN’, #balance.edge.weights = TRUE, metric=’angular’, score.component.variance=TRUE, verbose=TRUE). Cell cluster communities and graph embeddings were processed with default settings, prior to label propagation with defaults con$propagateLabels. Labels were then back propagated to the Seurat object metadata for comparisons against initial cluster annotations and downstream processing.

#### Neural Net Cell Type Predictions

Cell types were predicted using a neural net algorithm through the scCAMEL package, as previously described for hDRG (Hu et al., 2022; Yu et al., 2024). Processed RNA-seq data was extracted from the relevant Seurat objects, including previously published spatial-seq (Tavares-Ferreira et al., 2022) and a cross-species harmonized atlas (Bhuiyan et al., 2024). For the latter, human barcodes were extracted, and gene names were converted from mouse to human with the biomaRt package (R).

Count and metadata were reassembled into AnnData objects in python for processing. Large clusters were downsampled to the mean (cluster size) +0.5*std to minimize a size bias. Data were then filtered for low expression and cell cycle genes prior to normalization in line with the scCAMEL documentation. Hyperparameter tuning was performed with Optuna using the C2 scRNA-seq data as the test/train datasets (learningRate=0.01537, optimizerMmentum=0.8951, dropout=0.42, https://arxiv.org/abs/1907.10902). After tuning, cell type predictions were performed against lumbar (visium) as well as lumbar, thoracic, and TG (harmonized) datasets.

#### Interactome Analysis

Previously published curated ligand receptor database (Wangzhou et al., 2021), was intersected with the single-nucleus RNA-seq data to identify mechanisms by which DRG endothelial cells, fibroblasts, and myelinating Schwann cells may signal to neurons. Cells for each non-neuronal cell type and neuronal subpopulation were pseudobulked and the expression (in counts per million) of all genes for each cell type was determined. The database was used to identify ligand genes that are expressed in each non-neuronal cell type, and corresponding receptors expressed in DRG neuron populations. Genes were considered expressed in a cell type if they were above the 20th percentile among all genes, as this threshold conservatively differentiated non- and lowly-expressed genes from highly expressed genes in kernel density plots of gene expression generated for each cell type. Additionally, ligand genes were filtered to include only those which were differently expressed (padj < 0.1) and upregulated in samples from chronic pain donors. The resulting interactome table includes 236 interactions between the non-neuronal cell types and each individual neuron subtype. SankeyMATIC was used to visualize the unique ligand-receptor interactions, and the ComplexHeatmap v.2.20.0 R package was used to visualize the expression of ligand and receptor genes. R v4.4.2 was used for all calculations and interactome generation.

#### Figure Generation

Figures were compiled in Inkscape, GraphPad Prism 10 (GraphPad Software, Inc. San Diego, CA USA), RStudio v.2024.12.0 and Adobe Illustrator 2024.

### Data availability

Processed data will be publicly accessible on the SPARC portal.

## Acknowledgements

The authors thank the patients for their contribution to the study. Conflict of Interest Statement: T.J.P. is a co-founder and holds equity in 4E Therapeutics, NuvoNuro, PARMedics, and Nerveli. T.J.P. has received research grants from AbbVie, Eli Lilly, Grünenthal, Evommune, Hoba Therapeutics, and The National Institutes of Health. M.C. is the chief medical officer and holds equity in 4E Therapeutics.

